# Presynaptic contact and activity opposingly regulate postsynaptic dendrite outgrowth

**DOI:** 10.1101/2022.07.27.501752

**Authors:** Emily L. Heckman, Chris Q. Doe

## Abstract

The organization of neural circuits determines nervous system function. Variability can arise during neural circuit development (e.g. neurite morphology, axon/dendrite position). To ensure robust nervous system function, mechanisms must exist to accommodate variation in neurite positioning during circuit formation. Previously we developed a model system in the *Drosophila* ventral nerve cord to conditionally induce positional variability of a proprioceptive sensory axon terminal, and used this model to show that when we altered the presynaptic position of the sensory neuron, its major postsynaptic interneuron partner modified its dendritic arbor to match the presynaptic contact, resulting in functional synaptic input (Sales et al., 2019). Here we investigate the cellular mechanisms by which the interneuron dendrites detect and match variation in presynaptic partner location and input strength. We manipulate the presynaptic sensory neuron by (a) ablation; (b) silencing or activation; or (c) altering its location in the neuropil. From these experiments we conclude that there are two opposing mechanisms used to establish functional connectivity in the face of presynaptic variability: presynaptic contact stimulates dendrite outgrowth locally, whereas presynaptic activity inhibits postsynaptic dendrite outgrowth globally. These mechanisms are only active during an early larval critical period for structural plasticity. Collectively, our data provide new insights into dendrite development, identifying mechanisms that allow dendrites to flexibly and robustly respond to developmental variability in presynaptic location and input strength.

## Introduction

Neural circuit organization dictates circuit function, influencing behavior, cognition, and perception. While developmental programs in genetically identical animals produce similar final products, variability arises during circuit wiring to produce differences in cellular morphology, synaptic partnerships, and numbers of synapses between partners (Mohr et al., 2004; Chou et al., 2010; Caron et al., 2013; Linneweber et al., 2020; Churgin et al., 2021; Courgeon and Desplan, 2019; Couton et al., 2015; Goodman, 1978; Tobin et al., 2017; Witvliet et al., 2021). Such variability can arise innately due to stochastic processes (e.g. filopodial extension/retraction, lateral signaling, gene expression) (Özel et al., 2015; Troemel et al., 1999; Wernet et al., 2006) or when environmental factors impinge on development (e.g. rearing temperature, sensory experience) (Hubel et al., 1977; Kiral et al., 2021; Shatz and Stryker, 1978). Neural development must be flexible to account for these variations in order to generate robust circuit function.

If the location and strength of synaptic inputs can vary, how do surrounding neurons adapt such that they receive the right type and right amount of input? Dendrites are the major sites of synaptic input onto a neuron. Studies have shown that dendrites alter their morphology in response to varying levels of synaptic input (Ackerman et al., 2021; Takeo et al., 2021; Tripodi et al., 2008), and more recently we showed that dendrites can alter their position when a presynaptic partner is routed to an alternate neuropil location (Valdes-Aleman et al., 2021). Dendrites are clearly capable of structural plasticity, yet how they appropriately respond to a variable developmental landscape is unclear.

Here we investigate how and when developing dendrites accommodate wiring variation to ensure robust circuit connectivity. Leveraging strategies to manipulate presynaptic activity levels and presynaptic contact with postsynaptic dendrites, we find that there are two opposing mechanisms used to regulate robust partner matching: presynaptic contact promotes dendrite outgrowth locally, while presynaptic activity inhibits elongation of postsynaptic dendrites globally. These two strategies highlight the important role of presynaptic inputs in regulating the placement and subsequent elaboration of postsynaptic dendrites.

## Results

### dbd-Gal4 labels the dbd sensory neuron prior to formation of presynaptic contacts

To ensure robust circuit function, developing neurons must exhibit specificity and flexibility – specificity in synaptic partner choice, and flexibility to respond to variability in partner neuropil territory. The goal of our study was to determine the cellular mechanisms used by dendrites to compensate for variability in presynaptic axon placement and input strength to facilitate functional connectivity. To study these mechanisms in vivo, we used a model system consisting of the *Drosophila* larval dbd sensory neurons, and their postsynaptic partners, the A08a interneurons. The dbd sensory neurons form segmentally-repeated connections with A08a in the larval abdominal segments. A08a has two distinct dendritic domains: lateral and medial. All major inputs to A08a synapse with a single dendrite, either lateral or medial; the dbd sensory neuron synapses with the medial dendrite (Sales et al., 2019; Schneider-Mizell et al., 2016).

We sought to induce variability in the placement of the dbd axon to determine the extent to which its connectivity with A08a is stringently required at the medial dendrite. To do this we first needed genetic access to dbd prior to its interaction with A08a. We previously used *165-Gal4* (subsequently referred to as *dbd-Gal4*) to label the dbd sensory neuron (Sales et al., 2019; Valdes-Aleman et al., 2021), but the onset of dbd-Gal4 expression was not determined. Here we use dbd-Gal4 to drive expression of a myr::HA tag, and the 22C10 antibody to label all sensory neurons as a landmark. We observed the first expression of dbd-Gal4 in dbd neurons at embryonic stage 14. At this time, the axons have just entered the dorsal CNS as immature growth cones (Figure 1A-A’), and would not yet have contacted the more ventral neuropil domain occupied by the A08a dendrites. By stage 15, the dbd neurons formed anterior/posterior bilateral branches adjacent to the midline of the neuropil (Figure 1B-B’), followed by further elaboration to link adjacent segments in stage 17 (Figure 1C-C’). By this time, dbd axon terminals occupy a more ventral region of the neuropil where A08a medial dendrites are formed (Schrader and Merritt, 2000; Zlatic et al., 2003). These patterns of dbd projection into the CNS are schematized in Figure 1D.

**Figure 1.**
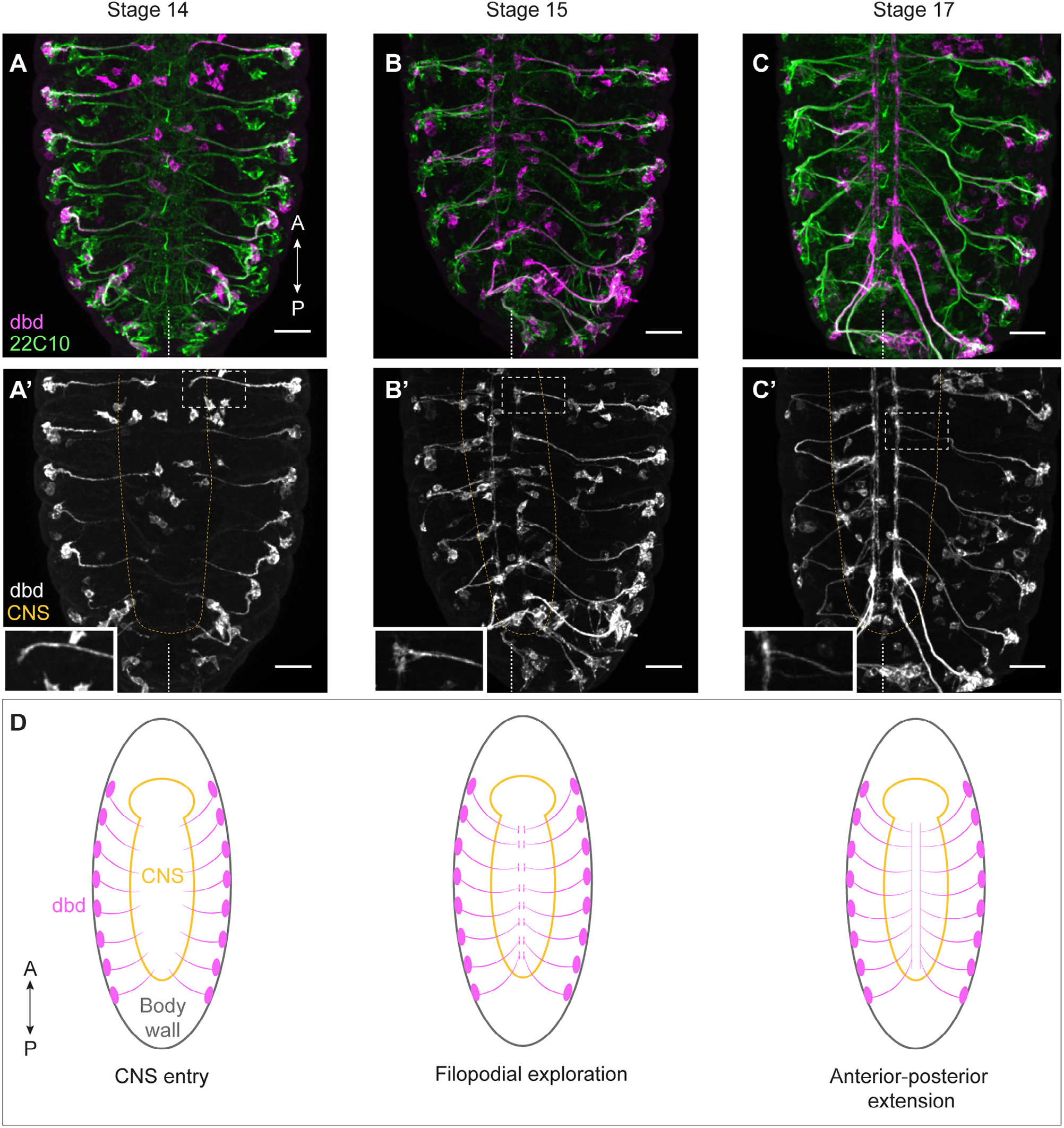
Onset of dbd-Gal4 expression during dbd axogenesis. (**A-C**) Stage 14, 15, and 17 fixed embryos. Sensory neurons labeled with 22C10 antibody (green) and dbd-Gal4 pattern labeled with smGdP-myr::HA (pink). Stage 14, n=4 animals; Stage 15, n=13 animals; Stage 17, n=17 animals. Pink cell bodies not in the body wall are likely part of the gut. A’-C’ show just the smGdP-myr::HA channel. Insets show zoomed in view of dbd outlined by white dashed box. Yellow dashed line indicates outline of central nervous system (CNS). Midline indicated by white dashed line at the bottom of each image. Scale bars, 20μm. (**D**) Illustrations summarizing results in A-C.

The A08a interneuron is labeled using a previously characterized LexA (*26F05-LexA*) to express *LexAop-myr::V5* in A08a and its dendritic arbors. This line first labels A08a in the early hours of the first larval instar (Figure 1 – figure supplement 1A-A’). By this time, A08a has contacted dbd and has developed its characteristic lateral and medial dendritic arbors. The timing of A08a-LexA expression precludes us from knowing the state of A08a dendrite development at the time of first contact with dbd. However, we conclude that dbd-Gal4 labels the dbd sensory neuron prior to its growth into the A08a neuropil domain, and thus it is an appropriate tool for manipulating dbd prior to the establishment of dbd-A08a synaptic connectivity.

### The dbd sensory neuron locally promotes dendrite elongation in the A08a interneuron

In a previous study, we tested the ability of dbd and A08a to compensate for developmentally-induced wiring variation (Sales et al., 2019). We tested for stringent specificity in dbd connectivity with the medial arbor by using genetic methods to target the dbd axon to the lateral neuropil. The dbd axon terminal could be misrouted to the intermediate and lateral neuropils through misexpression of repulsive axon guidance receptors, Robo-2 and Unc-5. dbd could form synapses at both intermediate and lateral dendritic arbors, and the strength of functional connectivity when dbd synapsed with the lateral arbor was indistinguishable from wild-type dbd-A08a connectivity (Sales et al., 2019).

Surprisingly, when dbd was targeted to lateral or intermediate neuropil domains, A08a produced an ectopic dendritic arbor to match the presynaptic contact (Valdes-Aleman et al., 2021). Here we extended and confirmed these findings by showing that the cumulative distribution of A08a dendrite volume corresponds to the location of dbd input (Figure 2A-D). The increase in dendrite arbor volume was most striking in the intermediate zone of the A08a dendritic domain where there are few arbors present in wild type (Figure 2A-B). Interestingly, when dbd was targeted to intermediate or lateral neuropil regions, the volume of the medial dendrite was decreased (Figure 2D).

**Figure 2.**
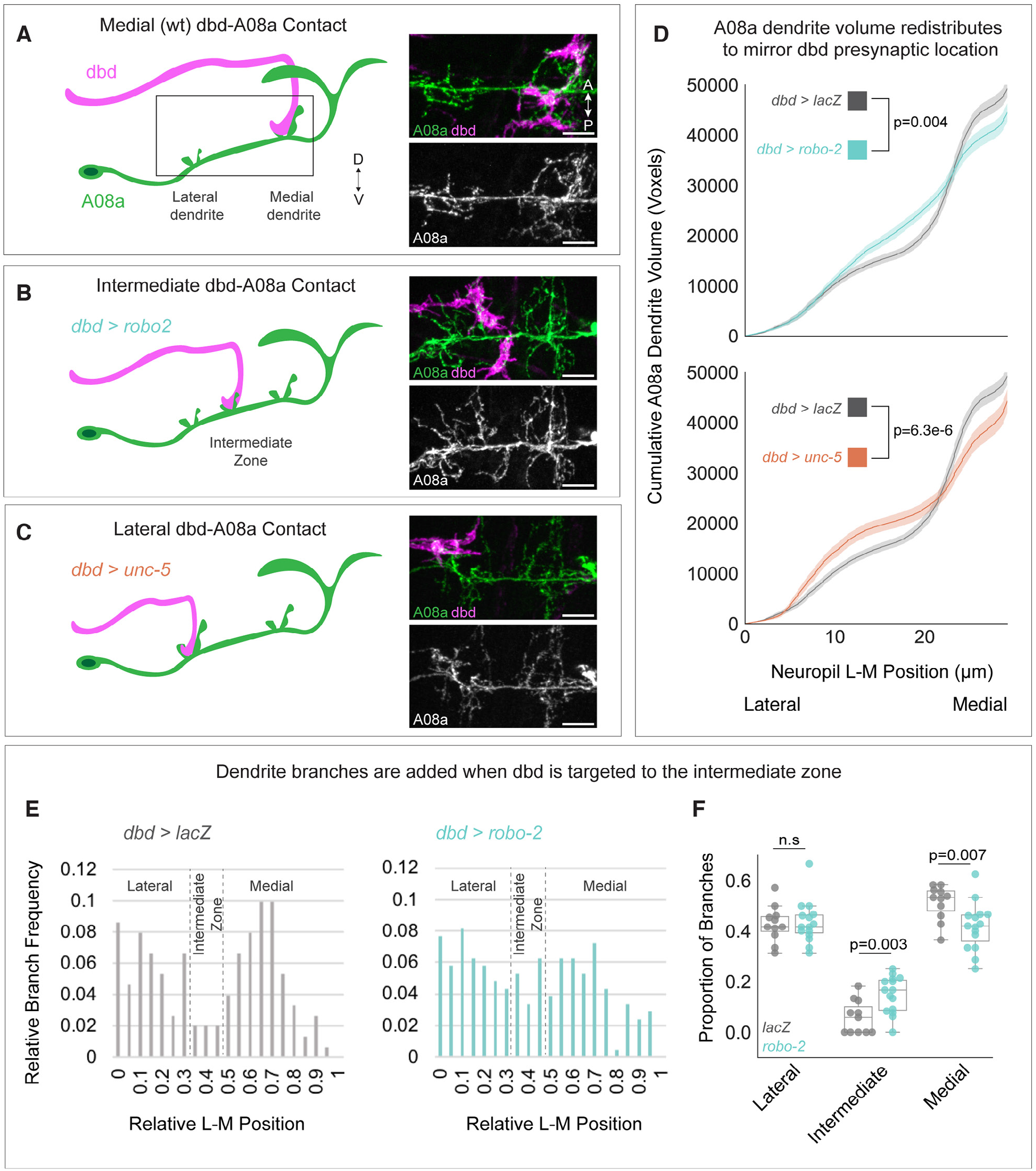
Dendrite development is promoted by presynaptic axons. (**A**) Left: Illustration of wild-type dbd (pink) and A08a (green). dbd projects to the A08a medial dendrite. Right: A08a dendritic domain (boxed region shown in cartoon). dbd (pink) contacts the medial A08a dendrite (green). Secondary image of A08a channel alone (white) shows two distinct dendritic domains. (**B**) Left: Robo2 misexpression leads dbd to project to the A08a intermediate dendritic domain. Right: dbd contacts the intermediate A08a dendritic domain, where there are ectopic dendrites. (**C**) Left: Unc-5 misexpression leads dbd to project to the A08a lateral dendrite. Right: dbd contacts the lateral A08a dendrite, where there is ectopic dendritic material. Scale bar, 5μm. Micrographs are from larvae aged 24 +/-4hrs alh. (**D**) Cumulative distribution of A08a dendrite volume (voxels) across the lateral-medial axis in conditions where dbd projects to the medial (gray, n=17 cells, 11 animals), intermediate (cyan, n=20 cells, 10 animals), or lateral (orange, n=11 cells, 10 animals) A08a dendrite. Solid line=mean distribution; shaded area=standard error of the mean (SEM). A08a dendrites receiving input from intermediate or lateral dbd neurons have significantly different volume distributions from wild-type dendrites (2-way Kolmogorov-Smirnov test). Note that the control LacZ trace is the same for top and bottom panels. (**E**) Relative Lateral-Medial distribution of A08a dendrite branch points from the main A08a neurite in conditions where dbd projects to the medial (gray, n=11 cells from 7 animals) or intermediate (cyan, n=15 cells from 8 animals) dendritic domain. Lateral, Intermediate, and Medial boundaries are demarcated based on the local minimum of the LacZ distribution. (**F**) Proportion of branches occupying Lateral, Intermediate, and Medial A08a dendritic domains when dbd projects to the medial dendrite (gray, n=11 cells from 7 animals) or intermediate zone (cyan, n=15 cells from 8 animals). Individual points represent single cells. When dbd projects to the intermediate domain, there are more A08a branches in the intermediate domain and fewer in the medial domain. Statistics computed using 2-tailed unpaired t-test with unequal variance.

The rearrangement of dendrite volume could be due to novel dendrite outgrowth from the main A08a neurite, or due to elaboration of pre-existing lateral or medial arbors. To distinguish between these possibilities, we plotted the frequency of branch points off of the main A08a neurite across the lateral-medial axis. We found that when dbd is targeted to the intermediate zone, there was an increase in the frequency of dendritic branches off the primary A08a neurite in the intermediate zone, supporting the idea that the observed increase in dendrite volume in the intermediate zone is due to novel dbd-promoted dendrite outgrowth (Figure 2E-F). We conclude that presynaptic contact can locally promote postsynaptic dendrite outgrowth, ultimately ensuring robust partner matching.

### dbd ablation results in A08a lateral dendrite expansion

Our findings led us to investigate the mechanisms of ectopic A08a dendrite establishment. To test the hypothesis that dbd contact promotes local dendrite outgrowth, we genetically ablated dbd using the *dbd-Gal4* line driving expression of the pro-apoptotic gene *hid*. If dbd promotes medial dendrite outgrowth, we would expect dbd ablation to reduce the A08a medial dendritic arbor.

We confirmed dbd ablation by 22C10 staining, which labels all sensory neurons. Embryonic and larval sensory neuron cell bodies are located in the body wall, and have stereotyped positions. 22C10 marks the dbd neuron cell body, positioned at the base of the dorsal-most cluster of sensory neurons (Figure 3 - figure supplement 1A,C) (Ghysen et al., 1986). In addition, we also used the absence of dbd-Gal4 driven myr::HA to identify segments where dbd was ablated (Figure 3C; Figure 3 - figure supplement 1D). Hid expression indeed led to a loss of dbd neurons, as detected through the absence of both 22C10^+^ and HA^+^ cell bodies from the body wall (Figure 3 - figure supplement 1B,D). After confirming that Hid expression eliminated dbd, we next assayed control and dbd ablation larvae at 24 +/-2hrs after larval hatching (alh) for A08a dendrite length. A08a dendrites were reconstructed using the Imaris software Filaments tool (Figure 3B’, D’). As expected, controls showed well-branched medial and lateral dendritic arbors by immunostaining (Figure 3A-B) and in the Imaris reconstructions (Figure 3B’). In contrast, dbd ablation led to qualitatively enlarged lateral dendrites by immunostaining (Figure 3C-D) and in the Imaris reconstructions (Figure 3D’). Quantification confirmed that complete dbd ablation led to A08a lateral dendrites that were longer and more branched (Figure 3E-F). In contrast, there was no significant change in medial dendrite length or branching when dbd was ablated (Figure 3E-F).

**Figure 3.**
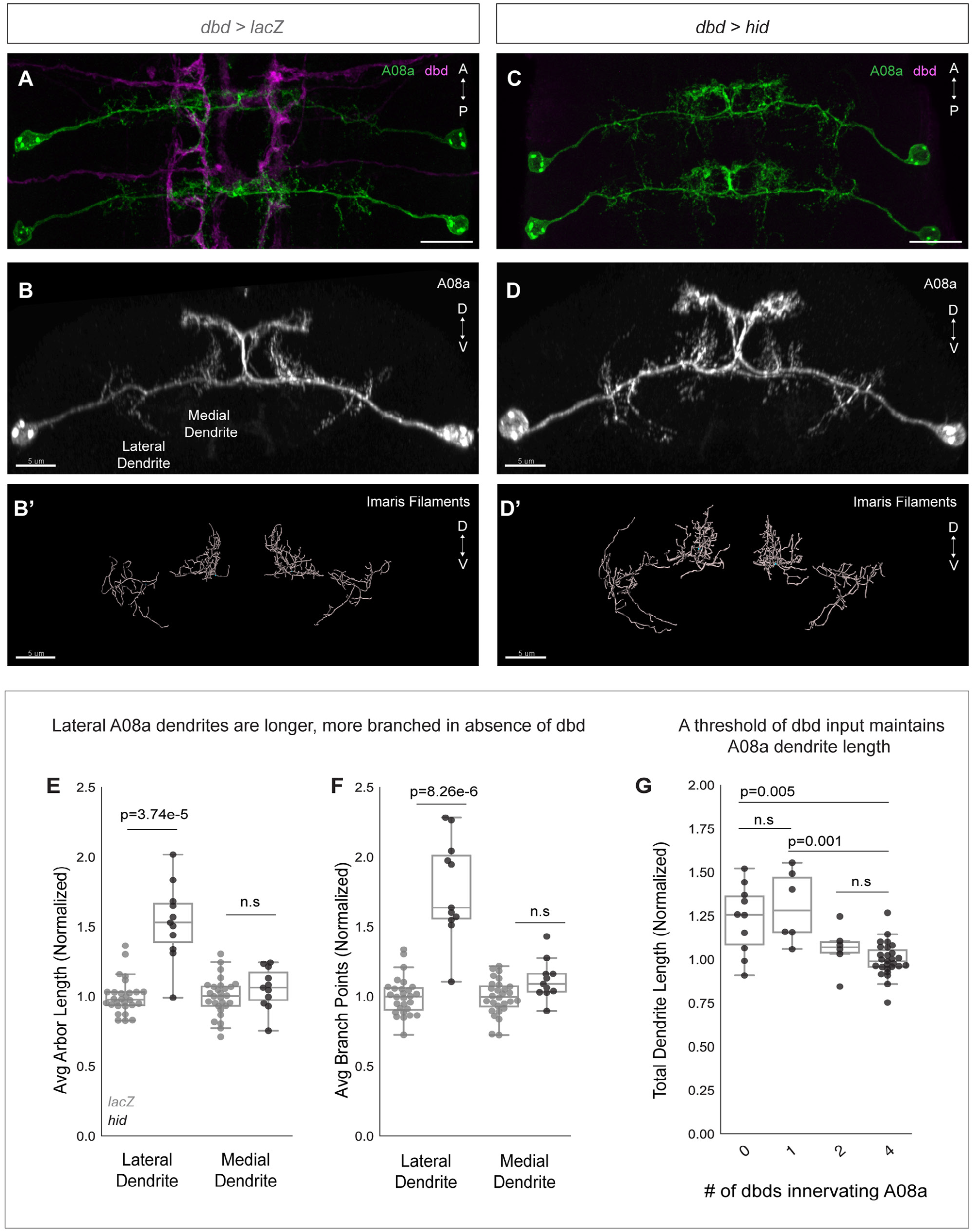
dbd ablation causes A08a lateral dendrite expansion. (**A**) Control VNC at 24 +/-2hrs alh, dorsal view. A08a neurons (green) are innervated by dbd neurons (magenta). Scale bar, 10μm. (**B**) Control A08a neurons from abdominal segment 1 of (A), posterior view. Scale bar, 5μm. (**B’**) Imaris filament reconstruction of A08a dendrites in (B). Scale bar, 5μm. (**C**) dbd ablation VNC at 24 +/-2hrs alh, dorsal view. A08a neurons (green) lack innervation from dbd neurons (magenta). (**D**) A08a neurons in dbd ablation background from abdominal segment 1 of (C), posterior view. Scale bar, 5μm. (D’) Imaris filament reconstruction of A08a dendrites in (E). Scale bar, 5μm. (**E**) Average dendrite length of lateral and medial A08a dendritic arbors in LacZ (control, gray, n=28 animals) or Hid-expressing animals (black, n=11 animals). (**F**) Number of dendrite branch points of lateral and medial A08a dendritic arbors in LacZ (control, gray, n=28 animals) or Hid-expressing animals (black, n=11 animals). Values normalized to control mean for either lateral or medial arbor. Circles represent single-animal averages between left and right hemisegments. Values for Hid-expressing animals with 0 remaining dbd neurons innervating A1 segment. (**G**) Total A08a dendrite length when A08a is innervated by 4 (LacZ control, n=28 animals), 2 (n=6 animals), 1 (n=6 animals), or 0 (n=10 animals) dbd neurons. Black circles represent single-animal dendrite length summed across left and right hemisegments. Data are normalized to dendrites innervated by 4 dbd neurons (LacZ control).Statistics computed using 2-tailed unpaired t-test with unequal variance.

Hid overexpression resulted in variable numbers of ablated dbd neurons across samples (Figure 3 - figure supplement 1B). We could therefore test whether there is a correlation between the number of dbd neurons innervating a segment and the total A08a dendrite length. In wild type, there are four dbd neurons innervating a single VNC segment, two per hemisegment. We found that when 1-2 of the four dbd neurons are ablated, A08a dendrite length is not significantly different than in controls. However, when 3-4 of the four dbd neurons are ablated, A08a lateral dendrite length is significantly increased (Figure 3G). This finding suggests that a threshold level rather than a linear summation of dbd input stabilizes A08a dendrite outgrowth.

The expansion of lateral A08a dendrite length was surprising, as dbd does not contact the lateral dendrite in wild-type animals (Sales et al., 2019; Schneider-Mizell et al., 2016). The growth in lateral dendrites following dbd ablation revealed that dbd provides negative regulation of A08a dendrite outgrowth, in addition to the positive mechanism revealed by the dbd lateralization experiments. Moreover, the negative mechanism acts throughout the neuron, rather than locally as does the positive mechanism. A likely source of a neuron-wide inhibitor of dendrite outgrowth is neuronal activity, which negatively regulates dendrite arbor size in multiple systems (Ackerman et al., 2021; Shen et al., 2020; Tripodi et al., 2008; Wu and Cline, 1998). We address this possibility in the next section.

### dbd activity globally inhibits A08a dendrite outgrowth

To test if dbd activity influences A08a dendrite size, we used two methods to silence dbd synaptic activity throughout development. The first was dbd-specific expression of either tetanus toxin light chain (TNT) or mutationally inactive TNT as a negative control. TNT cleaves the synaptic vesicle protein synaptobrevin, inhibiting evoked synaptic vesicle release (Sweeney et al., 1995). A previous study showed that dbd silencing led to slowed and uncoordinated larval locomotion (Hughes and Thomas, 2007). We validated that our silencing tools were effectively inhibiting dbd activity by comparing larval crawling behavior between control and dbd-silenced animals. Dbd-silenced animals had fewer waves of crawling activity, consistent with previous results (Figure 4A-B, E). Thus, we proceeded to ask if dbd silencing leads to expanded A08a dendrites. We found that constitutive dbd silencing using TNT indeed resulted in longer, more branched lateral and medial A08a dendrites at 26 +/-2hrs alh (Figure 4F-I).

**Figure 4.**
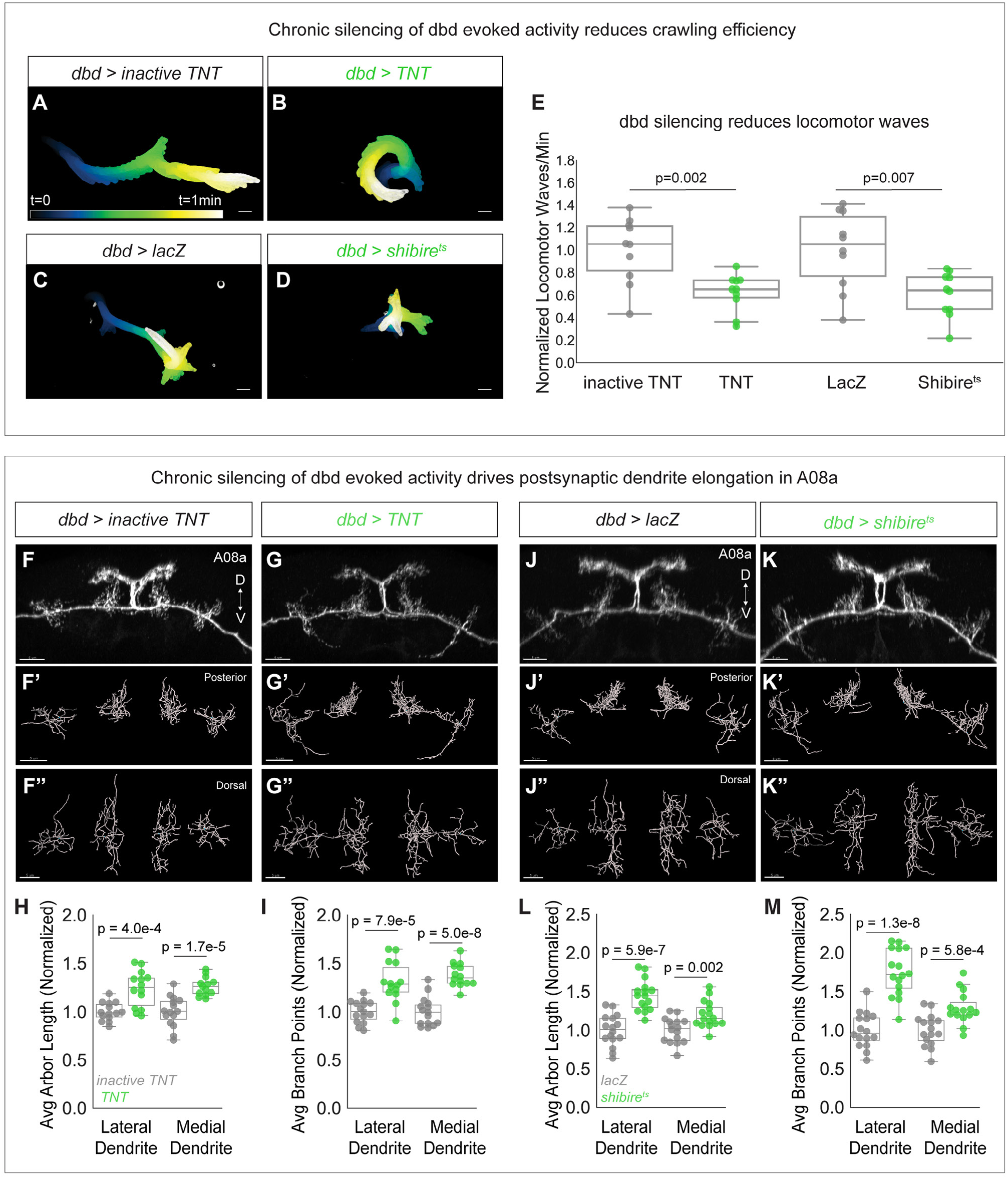
Chronic silencing of dbd activity drives A08a dendrite elongation. (**A**) Representative crawling trace of control inactive TNT control larva (26 +/-2hrs alh). Trace is color-coded by time. Scale bar, 1mm. (**B**) Representative crawling trace of TNT larva (26 +/-2hrs alh). (**C**) Representative crawling trace of control LacZ-expressing larva (24 +/-2hrs alh). Trace is color-coded by time. Scale bar, 1mm. (**D**) Representative crawling trace of Shibire^ts^ larva (24 +/-2hrs alh). (**E**) Number of locomotor waves (forward and reverse) initiated in one minute, normalized to corresponding control. Control animals (gray) initiate more locomotor waves relative to animals with silenced dbd neurons (green). inactive TNT, n= 10animals ; TNT, n= 10animals; LacZ, n=10 animals; Shibire^ts^, n= 10 animals. Statistics computed using 2-tailed unpaired t-test with unequal variance. Circles represent locomotor waves of single animals. (**F**) Control A08a neurons at 26 +/-2hrs alh, posterior view. (F’) Posterior view of Imaris filament reconstruction of A08a dendrites in (F). (**F”**) Dorsal view of (**F’**). (G) A08a neurons receiving input from TNT-expressing dbds at 26 +/-2hrs alh, posterior view. (**G’-G”**) Imaris Filament reconstructions of dendrites in (B). (**H**) Average dendrite length of lateral and medial A08a dendritic arbors in inactive TNT (control, gray, n=16 animals) or TNT-expressing animals (green, n=19 animals). (**I**) Number of dendrite branch points of lateral and medial A08a dendritic arbors in inactive TNT (control, gray, n=16 animals) or TNT-expressing animals (green, n=19 animals). (J) Control A08a neurons at 24 +/-2hrs alh, posterior view. (**J’-J”**) Imaris Filament reconstructions of dendrites in (J). (**K**) A08a neurons from receiving input from Shibire^ts^-expressing dbd, posterior view. (**K’-K”**) Imaris Filament reconstructions of dendrites in (K). (**L**) Average dendrite length of lateral and medial A08a dendritic arbors in LacZ (control, gray, n=14 animals) or Shibire^ts^-expressing animals (green, n=19 animals). (**M**) Number of dendrite branch points of lateral and medial A08a dendritic arbors in LacZ (control, gray, n=14 animals) or Shibire^ts^-expressing animals (green, n=19 animals).

We also used Shibire^ts^ to chronically and specifically silence dbd (Kitamoto, 2001). Shibire^ts^ animals were reared constitutively at 30°C and compared to temperature-matched negative controls expressing LacZ, as developmental temperature was recently shown to impact the extent of neurite branching and synapse formation (Kiral et al., 2021). Silencing of dbd with Shibire^ts^ also impaired larval crawling efficiency (Figure 4C-E), and resulted in longer, more branched A08a dendrites (Figure 4J-M). Presynaptic activity from dbd is therefore necessary to prevent excessive postsynaptic dendrite outgrowth in A08a.

If presynaptic activity inhibits A08a dendrite outgrowth, we predicted that elevated levels of dbd activity would result in shorter A08a dendrites. To activate dbd, we expressed the light-sensitive channelrhodopsin CsChrimson using dbd-Gal4 (Klapoetke et al., 2014). Animals were exposed to broad spectrum light throughout development and A08a dendrite length and complexity were assayed at 25 +/-2hrs alh. Compared to negative controls expressing LacZ, dbd optogenetic activation led to a decrease in overall A08a dendrite length and branching. This effect was most pronounced at the medial dendrite; although some lateral arbors exhibited decreased length and branching, it did not reach statistical significance (Figure 5A-D). The decreases in arbor length and branching were likely not due to excitotoxicity, as this method has been previously published in larval motor neurons without inducing excitotoxicity (Ackerman et al, 2021). We conclude that presynaptic activity is necessary and sufficient to restrict A08a dendrite outgrowth.

**Figure 5.**
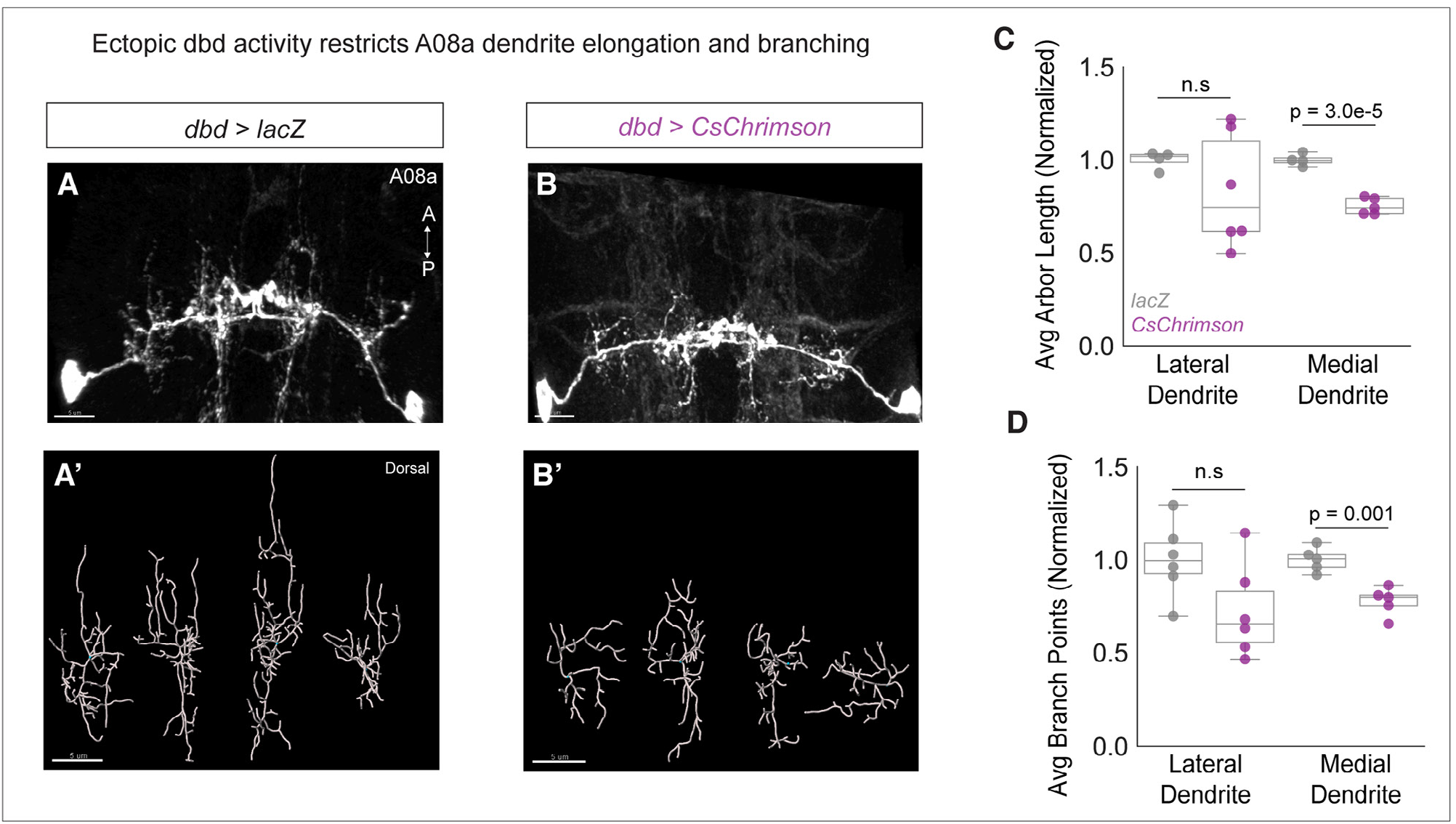
Chronic activation of dbd reduces A08a dendrite length. (**A**) Control A08a neurons control animal at 25 +/-2hrs alh, dorsal view. (**A’**) Imaris filament reconstructions of dendrites in (A). (**B**) A08a neurons receiving input from CsChrimson-expressing dbd neurons, activated throughout development. (**B’**) Imaris filament reconstructions of dendrites in (B). (**C**) Average dendrite length of lateral and medial A08a dendritic arbors in control (gray, n=4 animals) or CsChrimson-expressing animals (magenta, n=5-6 animals). (**D**) Number of dendrite branch points of lateral and medial A08a dendritic arbors in control (gray, n=5-6 animals) or CsChrimson-expressing animals (magenta, n=5-6 animals). Scale bars, 5μm. Values for all quantification normalized to control mean for either lateral or medial arbor. Circles represent single-animal averages between left and right hemisegments. Statistics computed using 2-tailed unpaired t-test with unequal variance.

### A08a dendrite plasticity is confined to a critical period of development

Neurons in many animals exhibit transient structural plasticity in early developmental windows that are termed “critical periods” for plasticity (Ackerman et al., 2021; Jarecki and Keshishian, 1995; LeVay et al., 1980; McLaughlin et al., 2003). In *Drosophila* larvae, motor neuron dendrites remain plastic until 8hrs alh, after which astrocytes prevent further dendritic remodeling, at least through 22hrs alh (Ackerman et al., 2021). Larvae also undergo continuous neuronal arbor growth to scale with their increasing body size, which may require some neurons to remain adaptable to a changing cellular environment (Gerhard et al., 2017). We therefore wanted to test whether A08a dendrites remain plastic in later stages of larval life, or if they are subject to the same critical period as motor neurons. To do so, dbd neurons were conditionally ablated by expressing Hid at successive stages of development, and A08a dendrite length and branching were quantified (Figure 6A).

**Figure 6.**
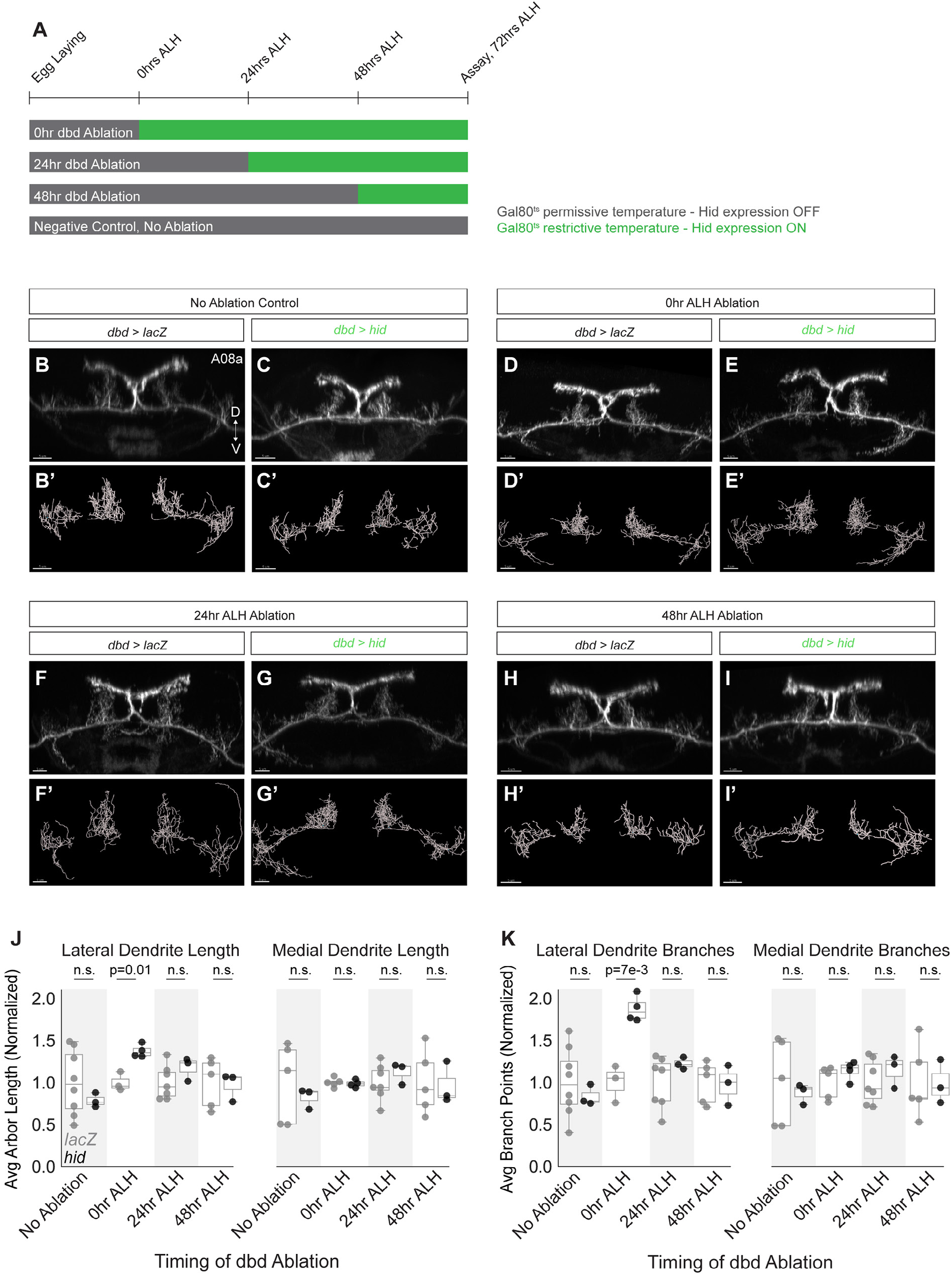
A08a dendrite plasticity is confined to a critical period in larval development. (**A**) Experimental design. Hid expression was inhibited at 18°C by Gal80^ts^ (gray bars). Hid expression was induced by shifting animals to 30°C (green bars) at the 25°C equivalent of 0hrs alh, 24hrs alh, and 48hrs alh. A08a dendrites were assayed for length and branching at the 25°C equivalent of 72hrs alh. (**B**) Control A08a raised continuously at 18°C. Imaris filament reconstructions of A08a dendrites in B’. (**C**) A08a in hid-expressing animal raised continuously at 18°C. Imaris filament reconstructions of A08a dendrites in (C’). (**D**) A08a in control animal shifted to 30°C at 0hrs alh. Imaris filament reconstructions of A08a dendrites in (D’). (E) A08a in hid-expressing animal shifted to 30°C at 0hrs alh. Imaris filament reconstructions of A08a dendrites in (E’). (**F**) A08a in control animal shifted to 30°C at 24hrs alh. Imaris filament reconstructions of A08a dendrites in (F’). (**G**) A08a in hid-expressing animal shifted to 30°C at 24hrs alh. Imaris filament reconstructions of A08a dendrites in (G’). (**H**) A08a in control animal shifted to 30°C at 48hrs alh. Imaris filament reconstructions of A08a dendrites in H’. (**I**) A08a in hid-expressing animal shifted to 30°C at 48hrs alh. Imaris filament reconstructions of A08a dendrites in (I’). (**J**) Average dendrite length of lateral (left) and medial (right) A08a dendritic arbors in LacZ (control, gray) or Hid-expressing animals (black). (**K**) Average number of branch points on lateral (left) and medial (right) A08a dendrites. X-axes, timing of dbd ablation (No ablation control: control n=5-8 animals, Hid n= 3 animals; 0hr alh ablation: control n=3-5 animals, Hid n=4 animals; 24hr alh ablation: control n=7 animals, Hid n=3 animals; 48hr alh ablation: control n=5 animals, Hid n=3 animals). For Hid quantifications, only segments containing 0-1 dbds were analyzed (determined by absence of dbd membrane stain). Scale bars, 5μm. Values for all quantification normalized to control mean for each ablation timepoint. Circles represent single-animal averages between left and right hemisegments. Statistics computed using 2-tailed unpaired t-test with unequal variance.

We controlled the onset of Hid expression in dbd using temperature-sensitive Gal80 (Gal80^ts^). Hid expression was induced at 0hrs alh, 24hrs alh, or 48hrs alh (times adjusted to 25°C developmental equivalent). A08a dendrite length was assayed for all experiments at 72hrs alh (Figure 6A). If A08a retains the capacity for dendrite plasticity throughout larval life, we would detect increases in dendrite length and branching after ablating dbd at each timepoint. In contrast, compared to no ablation controls (Figure 6B,D,F,H,J,K), we found that A08a was competent to expand its lateral dendrites only after dbd was ablated in newly hatched larvae (Figure 6D-E’, J-K), but not when ablations occurred at 24 or 48hrs alh (Figure 6F-K). This result demonstrates that A08a dendrite structural plasticity is confined to an early critical period, perhaps the same critical period as used by the *Drosophila* larval motor system (Ackerman et al., 2021).

## Discussion

### Presynaptic contact promotes local dendrite outgrowth

We sought to uncover the cellular mechanism by which dendrites respond to variable positioning and input of their synaptic partners. Here and in our previous work, we rerouted the dbd axon terminal to the lateral and intermediate neuropils and found that A08a dendrites mirrored the location of their displaced presynaptic partner (Valdes-Aleman et al., 2021). When dbd was targeted to the A08a intermediate dendritic domain, an area devoid of dendrites in wild-type, ectopic branches were established. At the same time, we observed a decrease in medial dendrite volume and branching when dbd was targeted elsewhere. In these experiments we measured no significant change to dbd-A08a functional connectivity strength (Sales et al., 2019), indicating that these compensations in dendrite length were likely activity-independent and due to contact alone.

Across a variety of model systems, presynaptic contact is correlated with or promotes the local outgrowth of dendrites (Chen et al., 2010; Jacoby and Kimmel, 1982; Kamiyama et al., 2015; Niell et al., 2004; Vaughn, 1989). In classic studies performed on the giant Mauthner (M) cells in zebrafish and axolotl, ablation of sensory afferents resulted in failed M-cell dendrite formation, whereas suprainnervation by these afferents was sufficient to cause overelaboration of the M-cell dendrites (Goodman and Model, 1988; Kimmel et al., 1981, 1977). Interestingly, when paired with a pharmacological silencing manipulation, suprainnervation of the M-cell still resulted in elongated dendrites, suggesting that contact-based cues are sufficient to drive local dendrite outgrowth (Goodman and Model, 1990), matching our results. Our similar results when dbd is mistargeted suggest a conserved mechanism across vertebrates and invertebrates for the local initiation of dendrites by sensory afferents. The ability for an axon to promote local dendrite outgrowth offers a potential strategy for pre- and postsynaptic partner matching that is robust to variable axon positioning.

One outstanding question from our studies is whether the dbd axon induces A08a dendrite formation de novo, or selectively stabilizes nascent dendrites. There are currently no methods for tracking the initial formation of A08a dendrites. The LexA driver used to label A08a is first expressed in early larval life, after first outgrowth of A08a medial and lateral dendrites (Figure 1 – figure supplement 1A). At the time of dbd CNS innervation at embryonic stage 14, the morphology of A08a dendrites is unknown. Therefore it is unclear if dbd induces novel dendrite outgrowth from an otherwise bare neurite, or if it stabilizes and promotes continued dendrite elongation from an arbor already beginning to take form. In wild-type animals, A08a dendrites are innervated by multiple neurons (Sales et al., 2019). If dbd is not the first neuron to contact the A08a medial dendritic domain, it is possible that the normal function of the dbd axon is to promote the continuous elongation of the medial dendrite rather than its initial induction. Either possibility would support our finding that presynaptic contact promotes postsynaptic dendrite outgrowth, and clarifying the exact mechanism will be important for identifying the molecular players that support either process.

### Presynaptic contact and presynaptic activity have opposing effects on dendrite outgrowth

We further tested the hypothesis that presynaptic contact promotes local dendrite outgrowth by genetically ablating dbd. When dbd was ablated, A08a lateral dendrites were elongated while medial dendrites were unchanged. Direct silencing of dbd caused both lateral and medial dendrites to elongate; this result highlights that dbd also acts as a negative regulator of postsynaptic dendrite outgrowth, and that presynaptic activity can downregulate global dendrite development.

We hypothesize that when dbd is ablated, A08a medial dendrites are unchanged due to the opposing roles of presynaptic activity and contact. Lack of dbd contact (which restricts growth) and lack of dbd activity (which promotes growth) could be working in direct opposition and thus result in no change in medial arbor size. In contrast, the lateral arbor experiences only the loss of activity, leading to arbor growth. Taken together, our data support a model of dendrite development in which presynaptic contact acts locally to promote dendrite outgrowth, whereas presynaptic activity acts globally across an entire neuron to downregulate dendrite elongation (Figure 7). As synapses are added and become functional, activity could act as a negative-feedback mechanism to excessive dendrite elongation.

**Figure 7.**
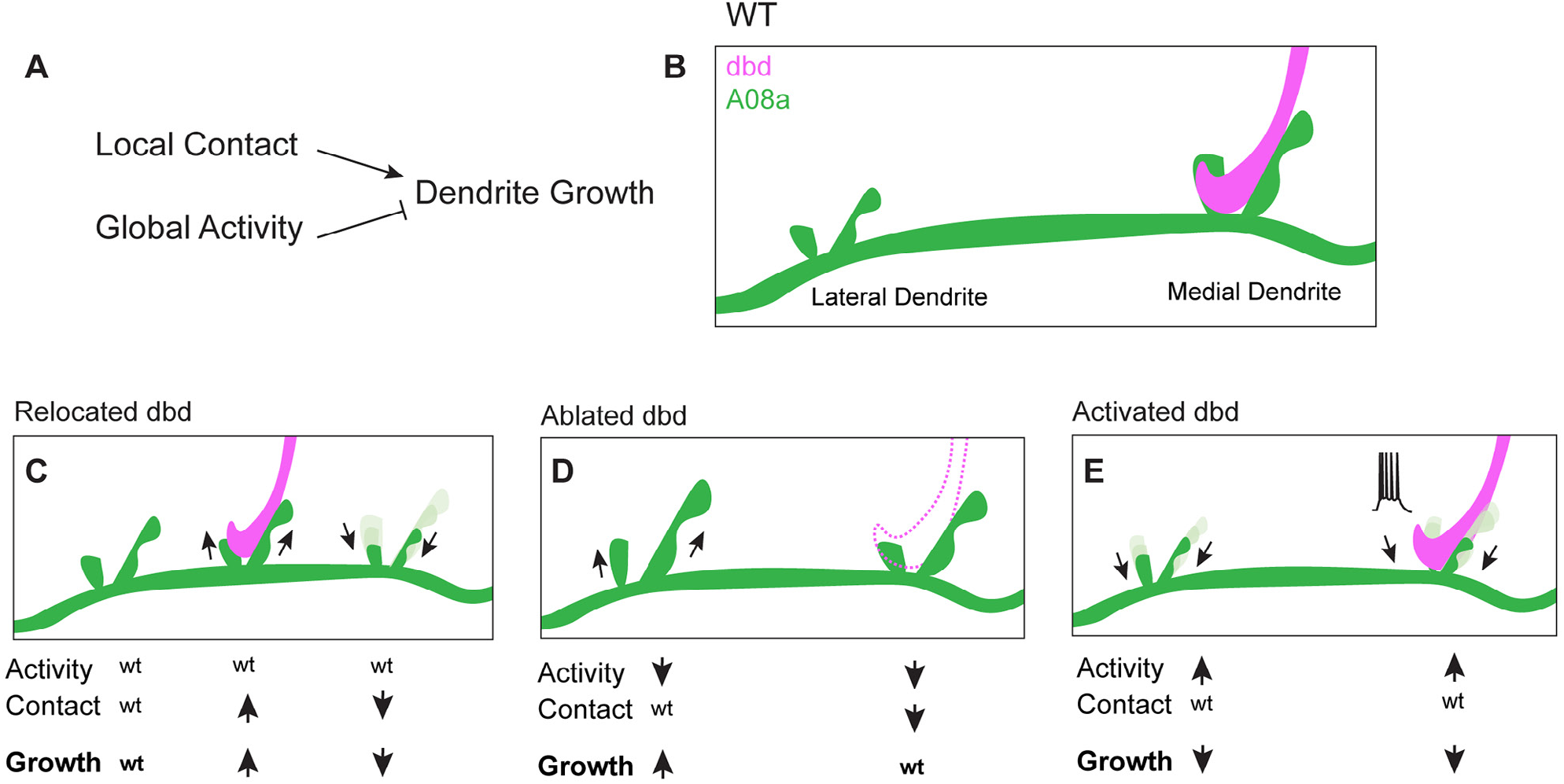
Proposed model: Presynaptic activity and contact opposingly regulate dendrite outgrowth. (**A**) Presynaptic contact promotes local dendrite outgrowth, while presynaptic activity levels inhibit neuron-wide dendrite outgrowth. (**B**) Wild type (WT) A08a dendrites. (**C**) When dbd is mistargeted to intermediate A08a dendritic domain, activity levels are wt. Local outgrowth is promoted at the intermediate dendrite, and inhibited by lack of contact at the medial dendrite. (**D**) When dbd is ablated, neuron-wide activity levels are decreased. This promotes lateral dendrite outgrowth. Lack of contact at the medial dendrite opposes the activity-dependent drive to elongate so dendrite length remains wt. (**E**) dbd Activation promotes neuron wide dendrite retraction/premature stabilization.

### Homeostatic vs. linear scaling of dendrite growth

Homeostatic structural plasticity is a phenomenon in which neurites adjust their length to counter the effect of too much or too little activity; when activity is excessive, the dendrite shrinks, and when activity is diminished, the dendrite expands. Homeostatic regulation of dendritic arbor size has been documented in insects (Ackerman et al., 2021; Hoy et al., 1985; Tripodi et al., 2008; Yuan et al., 2011) and vertebrates (Shen et al., 2020; Takeo et al., 2021; Tanvir et al., 2021; Wu and Cline, 1998). We observed a homeostatic relationship between presynaptic activity levels and postsynaptic A08a dendrite length. When synaptic input onto A08a was decreased by silencing evoked dbd activity, A08a dendrites were elongated; when dbd was chronically activated, A08a dendrites were smaller. Compensatory adjustments in dendrite length are likely a strategy to maintain a “set-point” of synaptic input. Such a mechanism would be useful to maintain a constant level of postsynaptic output when the amount of input is variable.

There are also examples in which synaptic activity positively correlates with dendrite elongation, a phenomenon we refer to as “linear scaling.” For example, studies in the *Xenopus* optic tectum showed that rearing tadpoles in the light (increased activity) can increase the rate of dendrite growth, and that growth is inhibited by pharmacologically blocking glutamate receptor activity (Rajan and Cline, 1998; Sin et al., 2002). In *Drosophila*, misexpression of activity-dependent transcription factors and ion channels can also promote dendrite elongation (Hartwig et al., 2008; Timmerman et al., 2013; Vonhoff et al., 2013). It remains an interesting open question why increased activity sometimes reduces dendrite size (homeostatic growth) and sometimes increases dendrite growth (linear scaling).

### A critical period for dendrite development in the *Drosophila* larva

A critical period for *Drosophila* larval motor neuron dendrite homeostasis was recently defined (Ackerman et al., 2021). In this system, dendrite length can be homeostatically modified by levels of activity. These motor dendrites lose the capacity to undergo activity-dependent remodeling at 8hrs alh, early in larval life. The closure of this critical period is governed by astrocytes - their infiltration into the neuropil coincides with critical period closure and their contact with motor dendrites prevents precocious dendrite extension/retraction (Ackerman et al., 2021; Stork et al., 2014).

From 1^st^ to 3^rd^ instar, the *Drosophila* larva increases in body size by two orders of magnitude. The nervous system continues to grow during this time, with individual neurons adding hundreds of microns of overall dendrite length (Gerhard et al., 2017). We wondered if an interneuron such as A08a would be subject to the same critical period described for motor neuron dendrites, or if dendrite plasticity remains necessary in subsequent stages of larval development to accommodate animal growth. We found that the capacity for A08a dendrites to respond to modified presynaptic input was confined to the first instar of larval development, similar to the critical period for motor dendrite structural plasticity (Ackerman et al., 2021). Ablation of dbd in subsequent larval instars did not impact A08a dendrite length. These results suggest that potentially all larval VNC neurons are subject to the same early critical period regulated by astrocytes, and structural plasticity events are not inducible in later stages of development. If presynaptic input is not required to regulate dendrite outgrowth in subsequent stages of larval life, perhaps separate cell-intrinsic (Tenedini et al., 2019; Zwart et al., 2013) or mechanical mechanisms (e.g. stretch or pulling forces) (Balice-Gordon and Lichtman, 1990; Bray, 1984; Tao et al., 2022) are required to allow the scaling of circuits as an organism grows.

### Dendrite diversification as a substrate for behavioral evolution

Here and in previous work, we observed that dendrite development is not hard-wired, but is modulated by the location of presynaptic partners and presynaptic activity levels (Sales et al., 2019; Valdes-Aleman et al., 2021). Within a species, subtle variation in neuron morphology that arises in development can diversify behavior. For example, natural variation in *D. melanogaster* Dorsal Cluster Neuron axonal projections directly impacts an animal’s ability to orient toward a visual object (Linneweber et al., 2020). Imprecise but constrained morphological development could be a “bet-hedging” strategy to promote the adaptability of an individual species to environmental changes.

Across species, it is intriguing to speculate that species-specific behaviors evolved in part through mechanisms that pattern neurite morphology. For two highly divergent nematode species, C. elegans and *P. pacificus*, amphid sensilla neuron number and soma position are highly conserved whereas ciliated dendrite morphology is more diverse. Correspondingly, neurons whose structure is more dissimilar between the species also have more divergent synaptic connections, implying that downstream behaviors would also differ (Hong et al., 2019). Another study found that in a species of *Drosophila* with evolved attraction to noni fruit, axon branch morphology in an olfactory processing center diverged from that of species that do not exhibit attraction to noni (Auer et al., 2020). For interneurons such as A08a, genomic changes resulting in alterations to dendrite neuropil position could vary the number and identities of presynaptic partners. It will be interesting to causally test the impact of neurite architecture on behavioral diversification, and specifically the extent to which genes regulating presynaptic axon position, neural activity, or critical period length are nodes of dendritic and ultimately behavioral evolution.

## Materials and Methods

### Key Resource Table

| Key Resources Table |  |  |  |  |
| --- | --- | --- | --- | --- |
| Reagent type (species) or resource | Designation | Source or reference | Identifiers | Additional information |
| Species (*Drosophila melanogaster*) | 26F05-LexA | BDSC | 54702 | Expressed in A08a neurons |
| Species (*D. melanogaster*) | 165-Gal4 | W. Grueber | N/A | Expressed in dbd neurons, as well as cIVda neurons |
| Species (*D. melanogaster*) | UAS-robo2::HA | BDSC | 66886 | Expresses HA-tagged Robo2 under UAS control |
| Species (*D. melanogaster*) | UAS-unc-5::HA | A. Zarin | N/A | Expresses HA-tagged Unc-5 under UAS control |
| Species (*D. melanogaster*) | UAS-hid, tubP-Gal80ts/CyO | M. Freeman | N/A | Expresses Hid under UAS control, Gal80ts negatively regulates Gal4 at 18°C. |
| Species (*D. melanogaster*) | UAS-TeTxLC-G2 | BDSC | 28838 | Expresses the light chain of tetanus toxin under UAS control |
| Species (*D. melanogaster*) | UAS-inactive TeTxLC | BDSC | 28840 | Expresses a mutated tetanus toxin light chain gene under UAS control |
| Species (*D. melanogaster*) | UAS-shi[ts1].K | BDSC | 44222 | Expresses temperature-sensitive shi under UAS control for inhibiting synaptic transmission at 30°C |
| Species (*D. melanogaster*) | UAS-LacZ | BDSC | 8529 | Control transgene |
| Species (*D. melanogaster*) | *UAS-LacZ* | BDSC | 8530 | Control transgene |
| Species (*D. melanogaster*) | *10xUAS-IVS-myr::smGdP::HA, 13xLexAop2-IVS-myr::smGdP::V5* | BDSC | 64092 | Expresses HA membrane tag under UAS control, V5 membrane tag under LexAop control |
| Species (*D. melanogaster*) | *20xUAS-IVS-CsChrimson.mVenus (attP2)* | BDSC | 55136 | Expresses CsChrimson tagged with mVenus under UAS control |
| Antibody, monoclonal | Mouse anti-V5 tag | Invitrogen, Carlsbad, CA, | Cat. R96025, Lot 1949337 | (1:1000) |
| Antibody, monoclonal | Rat anti-HA tag | Roche Holding, AG, Basel, Switzerland | Cat. 11867423001, Lot 27573500 | (1:100, after suggested dilution) |
| Antibody, monoclonal | Mouse anti-Futsch (22C10) | DSHB, Iowa City, IA | Concentrate 0.1mL | (1:1000) |
| Antibody, monoclonal | Rabbit anti-Corazonin | J. Veenstra, Univ Bordeaux | N/A | (1:2000) |
| Antibody, secondary | Alexa Fluor® 488 AffiniPure Donkey Anti-Mouse IgG (H+L) | Jackson ImmunoResearch, West Grove, PA | Cat. 715-545-151 | (1:400) |
| Antibody, secondary | Alexa Fluor® 647 AffiniPure Donkey Anti-Rat IgG (H+L) | Jackson ImmunoResearch, West Grove, PA | Cat. 712-605-153 | (1:400) |
| Antibody, secondary | Alexa Fluor® 488 AffiniPure Donkey Anti-Rat IgG (H+L) | Jackson ImmunoResearch, West Grove, PA | Cat. 712-545-153 | (1:400) |
| Antibody, secondary | Alexa Fluor® RRX AffiniPure Donkey Anti-Mouse IgG (H+L) | Jackson ImmunoResearch, West Grove, PA | Cat. 715-295-151 | (1:400) |
| Antibody, secondary | Alexa Fluor® 647 AffiniPure Donkey Anti-Mouse IgG (H+L) | Jackson ImmunoResearch, West Grove, PA | Cat. 715-605-151 | (1:400) |

### Fly stocks

| Genotypes | Figure |
| --- | --- |
| Females containing *10xUAS-IVS-myr::smGdP::HA*, *13xLexAop2-IVS-myr::smGdP::V5* (BDSC# 64092); *GMR26F05-LexA* (A08a neurons) (BDSC# 54702), *UAS-bruchpilot (short)-mstraw*; *165-Gal4* (dbd neurons) were crossed to males containing either (i) *UAS-lacZ* (BDSC #8529), (ii) *UAS-robo2::HA* (BDSC #66886), or (iii) *UAS-unc-5::HA* | (i) Figure 2A, D-F<br>(ii) Figure 2B, D-F<br>(iii) Figure 2C-D |
| Females containing *10xUAS-IVS-myr::smGdP::HA*, *13xLexAop2-IVS-myr::smGdP::V5* (BDSC# 64092); *GMR26F05-LexA* (A08a neurons) (BDSC# 54702), *UAS-bruchpilot (short)-mstraw*; *165-Gal4* (dbd neurons) were in-crossed to males of the same genotype. | Figure 1 A-C' |
| Females containing *10xUAS-IVS-myr::smGdP::HA*, *13xLexAop2-IVS-myr::smGdP::V5* (BDSC# 64092); *GMR26F05-LexA* (A08a neurons) (BDSC# 54702), *UAS-bruchpilot (short)-mstraw*; *165-Gal4* (dbd neurons) were crossed to males containing either (i) *UAS-lacZ.Exel* (control) (BDSC# 8529) or (ii) *UAS-hid, tubP-Gal80ts/CyO*. | (i) Figure 3A-B', E-G<br>Figure 6B, D, F, H, J-K<br>Figure 3 Sup 1B-C'<br>(ii) Figure 3C-D', E-G<br>Figure 6C, E, G, I, J-K<br>Figure 3 Sup 1B, D-D' |
| Females containing *10xUAS-IVS-myr::smGdP::HA*, *13xLexAop2-IVS-myr::smGdP::V5* (BDSC# 64092); *GMR26F05-LexA* (A08a neurons) (BDSC# 54702), *UAS-bruchpilot(short)-mstraw*; *165-Gal4* (dbd neurons) were crossed to males containing either (i) *UAS-inactive TeTxLC* (BDSC #28840) or (ii) *UAS-TeTxLC-G2* (BDSC #28838) | (i) Figure 4A, E-F", H-I<br>(ii) Figure 4B, E, G-I |
| Females containing *10xUAS-IVS-myr::smGdP::HA*, *13xLexAop2-IVS-myr::smGdP::V5* (BDSC# 64092); *GMR26F05-LexA* (A08a neurons) (BDSC# 54702), *UAS-bruchpilot(short)-mstraw*; *165-Gal4* (dbd neurons) were crossed to males containing either (i) *UAS-lacZ* (BDSC #8529) or (ii) *UAS-shi[ts1].K* (BDSC #44222) | (i) Figure 4C, E, J-J", L-M<br>(ii) Figure 4D-E, K-M |
| Females containing *10xUAS-IVS-myr::smGdP::HA*, *13xLexAop2-IVS-myr::smGdP::V5* (BDSC# 64092); *GMR26F05-LexA* (A08a neurons) (BDSC# 54702), *UAS-bruchpilot(short)-mstraw*; *165-Gal4* (dbd neurons) were crossed to males containing (i) *20xUAS-IVS-CsChrimson.mVenus* (attP2) (BDSC# 55136) or (ii) *UAS-lacZ* (BDSC #8530) | (i) Figure 5B-D<br>(ii) Figure 5A-A', C-D<br>Figure 1 Sup 1 |

### Animal Preparation

#### Embryo experiments

<u>Dbd-Gal4 Expression</u> (*Figure 1*): Embryos were collected overnight for 16hrs on 3.0% agar apple juice caps with yeast paste at 25°C. Embryonic stages were identified post-hoc by analyzing gut morphology. Stage 14, gut is tube shaped. Stage 15, gut is heart shaped. Stage 16, gut is coiled 3 times. Stage 17, gut is coiled 4 times.

<u>Hid Validation</u> (*Figure 3 – figure supplement 1*): Embryos were collected 4hrs on 3.0% agar apple juice caps with yeast paste at 25°C and were then aged at 30°C for 11hrs until approximately stage 17.

### A08a-LexA Expression (Figure 1 – figure supplement 1)

Embryos were collected on 3.0% agar apple juice caps with yeast paste for 4 hours at 25°C. Embryos were then aged for 21 hours. After 21hrs, embryos were transferred to a fresh 3.0% agar apple juice cap and then aged for 4 hours. Half of the hatched larvae were immediately dissected (aged 2 +/-2hrs ALH). The other half of the larvae were transferred to standard cornmeal fly food dishes and aged an additional 24hrs until dissection at 26 +/-2hrs alh. Due to stochastic expression of A08a-LexA in newly hatched larvae, samples were stained for Corazonin as a VNC segment landmark. Corazonin lables cells in T2-A6 (Choi et al., 2005).

### TNT Experiments (Figure 4)

Embryos were collected on 3.0% agar apple juice caps with yeast paste for 4 hours at 25°C. Embryos were then aged for 21 hours. After 21hrs, embryos were transferred to a fresh 3.0% agar apple juice cap and then aged for 4 hours. Hatched larvae were transferred to standard cornmeal fly food dishes and aged until dissection at 26 +/-2hrs alh.

### Shibire^ts^ and Hid experiments (Figure 3, Figure 3 – figure supplement 1, and Figure 4)

Embryos were collected on 3.0% agar apple juice caps with yeast paste for 4 hours at 25°C. Embryos were then aged for 17 hours at 30°C (Embryos and larvae develop 1.23x faster at 30°C). After 17hrs, embryos were transferred to a fresh 3.0% agar apple juice cap and then aged for 4 hours. Hatched larvae were transferred to standard cornmeal fly food dishes and aged at 30°C until dissection at 24 +/-2hrs alh.

### Chrimson Experiment (Figure 5)

All-*trans* retinal (ATR) is a necessary co-factor for CsChrimson. To ensure maternal transfer of ATR to larval progeny, parental crosses were fed yeast paste supplemented with ATR (final concentration 0.5mM; Sigma-Aldrich, R2500-100MG) for 72hrs. ATR yeast was made fresh daily and kept away from light. Embryos were then collected on 3.0% agar apple juice caps with +ATR yeast paste for 4 hours at 25°C. Embryos and larvae were aged continuously under broad spectrum light, approximately 10cm from the light source (∼30,000 lux; measured using Light Meter app for iPhone – Lightray Innovation GmbH). The temperature under the light was 28°C. Embryos age 1.1x faster at 28°C. After 19hrs, embryos were transferred to a fresh 3.0% agar apple juice cap and then aged for 4 hours. Newly hatched larvae were collected after the 4 hours and reared for an additional 23 hours under the light (larvae age 1.03x faster 28°C). Larvae were dissected at 25 +/-2hrs in low light (<100 lux) to prevent further Chrimson activation.

#### Critical Period Experiments (Figure 6)

Embryos were collected on 3.0% agar apple juice caps with yeast paste for 4 hours at 25°C.

#### No Ablation Group

Embryos were aged for 42hrs at 18°C (Embryos and larvae develop 2x slower at 18°C). After 42hrs, embryos were transferred to a fresh 3.0% agar apple juice cap and then aged for 4 hours. Hatched larvae were transferred to standard cornmeal fly food dishes and aged at 18°C until dissection at 146 +/-2hrs alh (25°C equivalent to 72hrs alh, middle of 3^rd^ instar).

#### 1^st^ Instar Ablation Group

Embryos were aged for 42hrs at 18°C. After 42hrs, embryos were transferred to a fresh 3.0% agar apple juice cap and then aged for 4 hours at 30°C. Hatched larvae were transferred to standard cornmeal fly food dishes and aged at 30°C until dissection at 67 +/-2hrs alh (25°C equivalent to 72hrs alh, middle of 3^rd^ instar).

#### 2^nd^ Instar Ablation Group

Embryos were aged for 42hrs at 18°C. After 42hrs, embryos were transferred to a fresh 3.0% agar apple juice cap and then aged for 4 hours. Hatched larvae were transferred to standard cornmeal fly food dishes and aged at 18°C for 48hrs. At this time animals were shifted to 30°C and raised an additional 44hrs until the time of dissection (25°C equivalent to 72hrs alh, middle of 3^rd^ instar).

#### 3^rd^ Instar Ablation Group

Embryos were aged for 42hrs at 18°C. After 42hrs, embryos were transferred to a fresh 3.0% agar apple juice cap and then aged for 4 hours. Hatched larvae were transferred to standard cornmeal fly food dishes and aged at 18°C for 96hrs. At this time animals were shifted to 30°C and raised an additional 22hrs until the time of dissection (25°C equivalent to 72hrs alh, middle of 3^rd^ instar).

No groups could be tested in which animals were reared continuously at 30°C until 3^rd^ instar as all animals expressing dbd>Hid died.

### Immunohistochemistry

#### Larval brain sample preparation

Larval brains were dissected in PBS, mounted on pre-EtOH treated 12mm #1thickness poly-D-lysine coated coverslips (Neuvitro Corporation, Vancouver, WA, Cat# GG-12-PDL) (primed in 70% EtOH at least one day prior to use). Samples fixed for 23 minutes in fresh 4% paraformaldehyde (PFA) (Electron Microscopy Sciences, Hatfield, PA, Cat. 15710) in PBST. Samples were washed in 0.3% PBST and then blocked with 2% normal donkey serum and 2% normal goat serum (Jackson ImmunoResearch Laboratories, Inc., West Grove, PA) in PBST overnight at 4°C or for 1hr at room temperature. Samples incubated in primary antibody for two days at 4°C. The primary was removed, and the samples were washed with 2 quick PBST rinses followed by 3×20min washes in PBST. Samples were then incubated in secondary antibodies overnight at 4°C, shielded from light. The secondary antibody was removed following overnight incubation and the brains were washed in PBST (2 quick rinses, followed by 3×20min washes). Samples were dehydrated with an ethanol series (30%, 50%, 75%, 100% ethanol; all v/v, 10 minutes each) (Decon Labs, Inc., King of Prussia, PA, Cat. 2716GEA) then incubated in xylene (Fisher Chemical, Eugene, OR, Cat. X5-1) for 2×10 minutes. Samples were mounted onto slides containing 2 drops of DPX mountant (Millipore Sigma, Burlington, MA, Cat. 06552) and cured for 1-3 days then stored at 4°C until imaged.

#### Embryo sample preparation

Embryos were transferred from apple caps into collection baskets and rinsed with dH_2_O. Embryos were dechorionated in 100% bleach (Clorox, Oakland, CA) for 3min and 30sec with gentle agitation. Dechorionated embryos were rinsed with dH_2_O for 1min. Embryos were fixed 25mins in 2mL Eppendorf tubes containing equal volumes of Heptane (Fisher Chemical, Eugene, OR, H3505K-4) and 4% PFA diluted in PBS. Fix was removed, and 850uL of Heptane was added to each tube. 650uL of Methanol were then added, and tubes were then subject to vigorous agitation for 1min in a step required for removing the vitelline membrane. Nearly all liquid was removed from the tubes, leaving the embryos. Embryos were rinsed in Methanol (Fisher Chemical, Eugene, OR, Lot# 206197, Cat. A412P-4) twice followed by 2 quick rinses in 0.3% PBST. PBST was removed and embryos were blocked with 2% normal donkey serum and 2 % normal goat serum (Jackson ImmunoResearch Laboratories, Inc., West Grove, PA) in PBST for 1hr at room temp. After blocking, embryos were incubated in primary and secondary antibodies and mounted as described above for larval brains.

### Light Microscopy

Fixed larval preparations were imaged with a Zeiss LSM 710 or LSM 900 laser scanning confocal (Carl Zeiss AG, Oberkochen, Germany) equipped with an Axio Imager.Z2 microscope. A 63x/ 1.40 NA Oil Plan-Apochromat DIC m27 objective lens and GaAsP photomultiplier tubes were used.

Software program used was Zen 2.3 (blue edition) (Carl Zeiss AG, Oberkochen, Germany). For each independent experiment, all samples were acquired using identical acquisition parameters.

### Image processing and analysis

#### Imaris Filament reconstruction and quantification of A08a dendrites (Figures 3-6)

Confocal image stacks were loaded into Imaris 9.5.1(Bitplane AG, Zurich, Switzerland). A08a dendrites from A1 and A2 segments were analyzed. A new Imaris Filament object was created for each A08a dendrite (lateral and medial). Briefly, the Filaments tool was selected, and a region of interest (ROI) drawn to encompass the dendrite. The source channel for A08a membrane (488) was selected. An approximation for minimum and maximum dendrite diameters were measured in Slice view, and found to be 0.2μm and 1μm respectively. These values were used to identify Starting Points and Seed Points for all images. Thresholds for Starting Points and Seed Points were manually adjusted until 1 Starting Point on the main A08a dendritic shaft remained, and Seed Points labeled A08a dendrite signal without labeling image background. The option to Remove Disconnected Seed Points was selected, with a Smoothing factor of 0.2μm. Absolute Intensity Threshold was manually adjusted until all Seed Points were filled. When the Filament was rendered, misidentified structures were selected and manually deleted or adjoined.

The sum length and the sum of all branch points of each dendritic arbor were calculated automatically in Imaris (Statistics > Details > Average Values). The values for left/right lateral dendrites and left/right medial dendrites were averaged for each animal.

For Figure 3G, the length of the A08a Left-Lateral Dendrite, Left-Medial Dendrite, Right-Medial Dendrite, Right-Lateral Dendrite was summed together, as left and right A08a’s form recurrent synaptic connections. The values were normalized to the mean total dendrite length of the WT controls (with 4 dbd inputs).

Sample exclusion criteria were established prior to conducting analysis. Samples were excluded from analysis if there was damage to the tissue or low signal-to-noise that obstructed the ability to reliably identify dendrite membrane signal.

#### A08a cumulative dendrite position (Figure 2D)

Published data from Sales et al., 2019 and Valdes-Aleman et al., 2021 were used. Larval brains aged 24 +/-4hrs alh were processed and imaged as described in Sales et al., 2019 and Valdes-Aleman et al., 2021. Briefly, image processing and analysis was performed using FIJI (ImageJ 1.50d, https://imagej.net/Fiji). Stepwise, images were rotated (Image > Transform > Rotate(bicubic)) to align dendrites of interest along the x axis, then a standardized ROI was selected in 3D to include the dendrites to analyze in one hemisegment (Rectangular selection > Image > Crop). To identify the voxels that contain dendrite intensity, a mask was manually applied (Image > Adjust > Threshold). The threshold was assigned to include dendrite positive voxels and minimize contribution from background. To quantify the amount of dendrite positive voxels across the medial-lateral axis, images were reduced in the Z-dimension (Image > Stacks > Z-project > Sum Slices) and a plot profile was obtained to measure the average voxel intensity (Rectangular selection > Analyze > Plot profile). The cumulative sum of A08a dendrite voxels was calculated for each individual hemisegment, and a mean voxel distribution was generated from the population data.

#### Branch distribution (Figure 2E-F)

Published data from Sales et al., 2019 and Valdes-Aleman et al., 2021 were used. Hemisegments from A1 and A2 were analyzed. Image analysis was performed in FIJI. Images were rotated to align the dendrites of interest along the x-axis (Image > Transform > Rotate(bicubic)). For each hemisegment analyzed, a rectangular ROI was drawn starting at the midline and ending at the lateral edge of the lateral-most branch point coming from the main A08a neurite. The width of the ROI was logged (in microns). The Cell Counter plugin was used to count the dendrite branches originating from the main A08a neurite (Plugins > Analyze > Cell Counter). The x position of each branch point was measured (in microns) (Cell Counter > Measure). The relative lateral-medial position of each branch point was determined by dividing the branch’s x position by ROI width. The relative frequency of branches at a given position was determined by counting the number of branches for that bin, and dividing that value by the total number of branches.

For Figure 2F, Lateral, Intermediate, and Medial domains were determined based on the relative peak branch positions in LacZ controls in Figure 1E. The Lateral domain = <0.35, Intermediate = ≥ 0.35 and <0.5, and Medial = >0.5. The proportion of branches in each domain was determined for each cell, and was then plotted as a single point in Figure 1F.

#### Figure preparation

Micrographs in figures were prepared as either 3D projections in Imaris 9.5.1 (Bitplane AG, Zurich, Switzerland) or maximum intensity projections in FIJI (ImageJ 1.50d, https://imagej.net/Fiji). Scale bars are given for reference on maximum intensity projections but do not necessarily represent actual distances, as the tissue samples undergo changes in size during the tissue clearing protocol. For images exported from the Imaris software, the scale bars are assigned to match the scale at the “center” of the 3D projection. Pixel brightness was adjusted in some images for better visualization; all adjustments were made uniformly over the entire image, and uniformly across corresponding control and experimental images. Larval crawling traces in Figure 3 – Supplement 1 are temporal projections made in FIJI (Image > Hyperstacks > Temporal Color-Code).

### Larval behavior assays

#### TNT experiments

Newly hatched larvae were aged for 24 hours on standard cornmeal fly food at 25°C. At this time, 5 larvae were gently transferred to a 4×4cm 1.2% agarose (Sigma, Lot# SLCD4639, Cat. A9539-500G) arena. Larvae were spaced apart to prevent collisions during recording. Prior to recording, larvae were acclimated to the arena for 2 minutes. The ambient temperature during recording was 20-22°C. Videos of individual larvae were collected at 5 frames/second for 1 minute.

#### Shibire experiments

Newly hatched larvae were aged for 22 hours on standard cornmeal fly food at 30°C. At this time, individual larvae were transferred to a 0.5mm thick 1.2% agarose pad positioned on top of a 22×40mm coverslip (Corning, Lot# 14418013, Cat. 2980-224, #1.5 thickness). The larva and coverslip were placed on top of a CherryTemp microfluidic chip (Cherry Biotech, Montreuil, France) with an 18°C surface temperature. Larvae were acclimated to the surface for 2 minutes prior to recording. Videos of individual larvae were collected at 5 frames/second for 1 minute. For all behavior experiments, animals that failed to move during the 1-minute recording were excluded from analysis.

#### Analysis of larval locomotor behavior

Movie processing and analysis was performed using FIJI (ImageJ 1.50d, https://imagej.net/Fiji). Locomotor waves were manually quantified. Forward and backward waves were summed together for each animal. The number of waves for each animal was normalized to the control average of each independent experiment.

##### Sample definition and in-laboratory replication

A minimum of two independent experiments were done for each experiment. Sample sizes were determined based on previously published standards. Data points in this paper describe biological replicates (multiple individual animals of a corresponding genotype), as technical replicates (repeated measures of the same sample) were not relevent.

### Statistical analyses

Statistics were computed using Excel or Python (scipy.stats). All statistical tests used are listed in the figure legends. For comparisons between cumulative distributions in Figure 2D, a 2-sample Kolmogorov-Smirnonv test was used as it is more appropriate for detecting differences across two distributions, rather than just looking for differences between the means of two groups. For remaining statistical tests, a student’s 2-way t-test was used to detect differences in the mean distribution. An assumption of unequal variance was used, as the sample sizes were not sufficiently large to assume equal variance. Statistical outliers were identified and exclulded by computing the 1^st^ and 3^rd^ quartiles for each group and setting upper and lower thresholds that deviated above or below the interquartile range (IQR) (lower threshold = 1^st^ Quartile – 1.5*IQR; upper threshold = 3^rd^ quartile + 1.5*IQR). P-values are reported in the figures. n.s. = not significant, where p>0.05. Plots were generated using Excel or Seaborn and Matplotlib packages in Python.

### Data availability

No new datasets were generated.

## Acknowledgements

We thank Dr. Bruce Bowerman and Dr. Molly Jud for the use of their CherryTemp system. We thank Alanna Sowles and Keiko Hirono for assistance with Imaris reconstructions. We thank Dr. Sarah Ackerman, Dr. Adam Miller, Dr. Cris Niell, and Peter Newstein for constructive comments on the manuscript. Stocks obtained from the Bloomington *Drosophila* Stock Center (NIH P40OD018537) were used in this study. Funding was provided by HHMI (CQD, ELH) and NIH HD27056 (CQD, ELH).

---

**Figure 1 – figure supplement 1.**
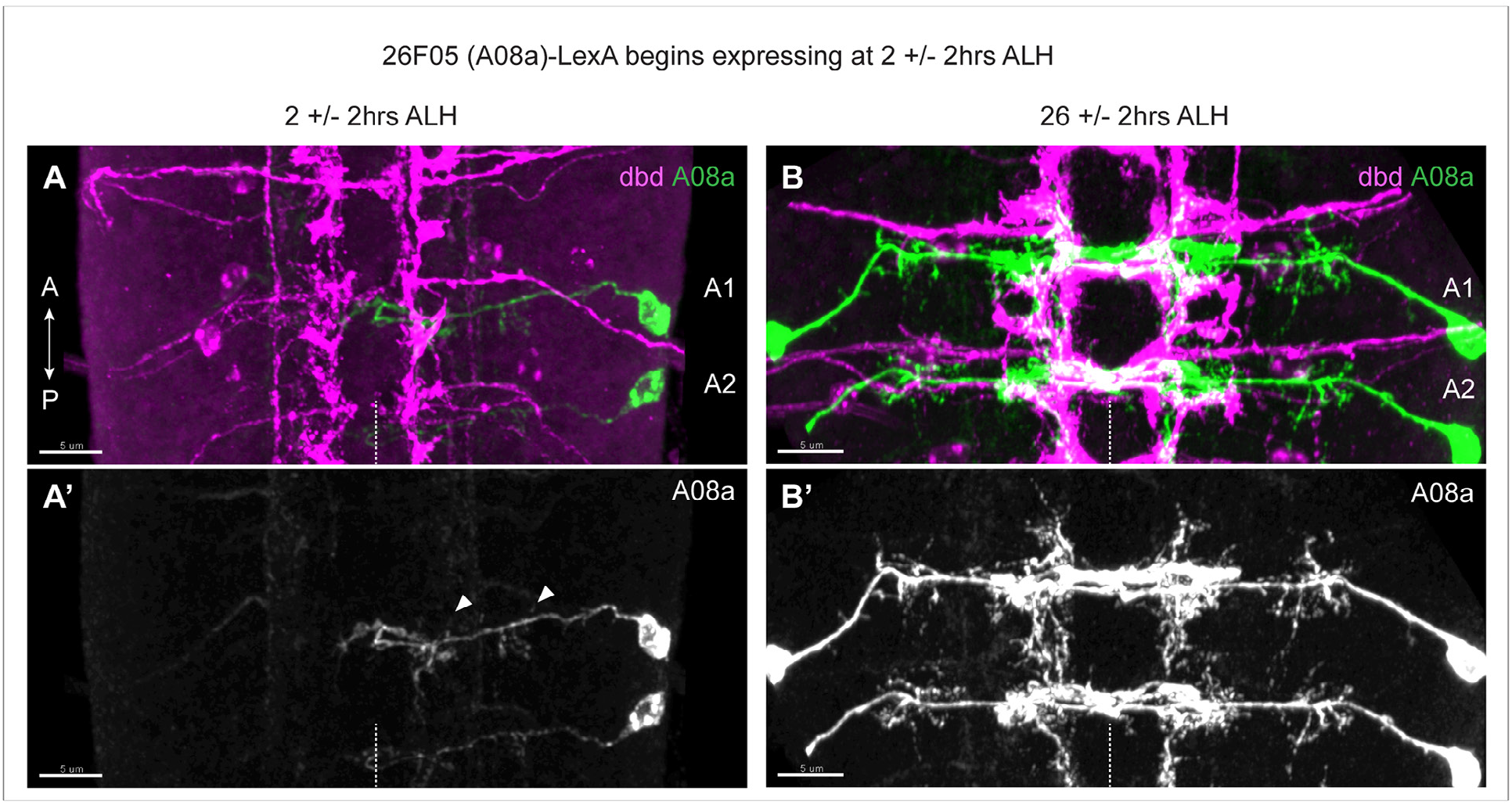
Onset of A08a-LexA expression during early larval life. (**A**) Dorsal view of VNC at 2 +/-2hrs ALH. Dbd-Gal4 expression pattern in pink, A08a-LexA expression pattern in green. (**A’**) A08a channel alone. Arrow heads indicate the medial and lateral dendrites. In this image, LexA expression is on in A1R, weakly in A2R, and not at all in the opposing A1L and A2L hemisegments. n=7 animals. (**B-B’**) Dorsal view of VNC at 26 +/-2hrs ALH. A08a-LexA is expressed robustly in all hemisegments at this time. n= 7 animals. For all images, scale bar = 5μm; midline indicated by white dotted line; and A1 and A2 label first and second abdominal segments.

**Figure 3 – figure supplement 1.**
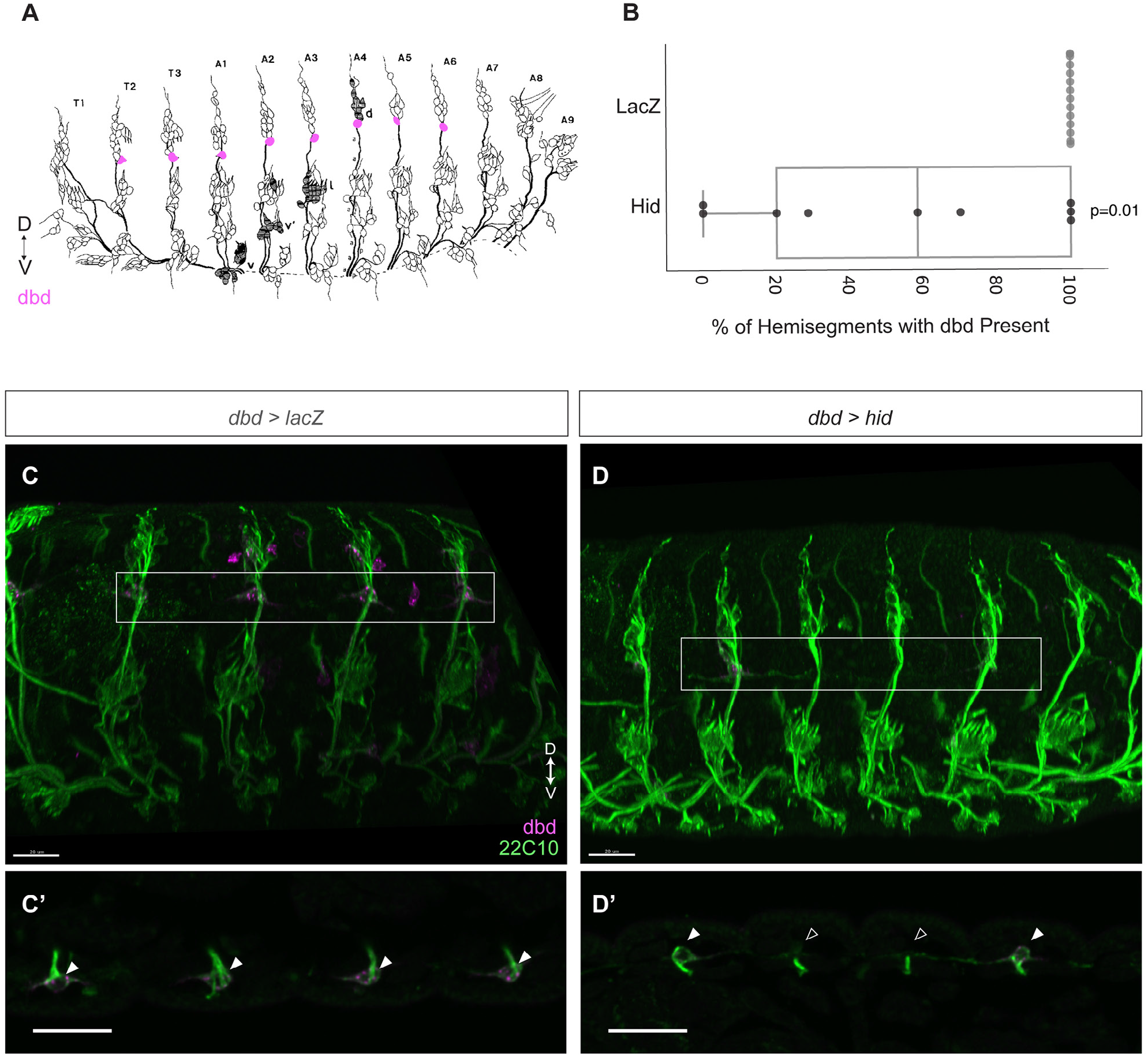
Validation of dbd ablation. (**A**) Illustration of embryonic sensory neuron anatomy (modified from Ghysen et al., 1986). Sensory neuron anatomy is segmentally repeated, and cell bodies are in stereotyped positions in the dorsal-ventral axis. Dbd cell bodies are highlighted in pink, positioned at the base of the dorsal-most cluster or cell bodies. Anterior is to the left. (**B**) Percentage of hemisegments containing an intact dbd cell body, identified by 22C10 staining in control (gray) or Hid-expressing (black) embryos. All control embryos had dbd 22C10 labeling in 100% of hemisegments (n=14 embryos). Most Hid-expressing embryos had missing dbd neurons (n=6 embryos). Hid embryos with 100% of dbd neurons intact (n=3 embryos) were likely CyO^+^ instead of Hid^+^ (see Methods). (**C-C’**) Stage 17 control embryo. 22C10 labels all sensory neurons (green), dbd-Gal4>myr::HA labels dbd and other cells (magenta). Scale bar, 20μm (C’) Substack of dbd cell bodies (filled arrowheads) boxed in (C). Scale bar, 20μm. (**D-D’**) Stage 17 Hid-expressing embryo. 22C10 labels all sensory neurons (green), dbd-Gal4>smGdP-myr::HA labels dbd and other cells. Scale bar, 20μm (D’) Substack of dbd cell bodies boxed in (D). dbd cell bodies are present in the first and last segment (filled arrowhead) and missing in the middle two segments (empty arrowhead), indicative of successful ablation. Scale bar, 20μm.

